# Coexistence of annual plants under competition and reproductive interference

**DOI:** 10.64898/2026.09.30.755560

**Authors:** Naoto Shinohara, Akira Yamawo, Koki R. Katsuhara

**Author notes:** Corresponding authors: Naoto Shinohara: (Email), Koki R. Katsuhara: (Email). These authors contributed equally (Naoto Shinohara, Koki R. Katsuhara).

## Abstract

In natural ecosystems, community dynamics are driven by diverse interactions operating through both ecological and reproductive processes. Although theory predicts that these processes jointly generate complex and contingent dynamics through the interplay of stabilizing (negative frequency dependence arising from niche differences) and destabilizing effects (positive frequency dependence caused by reproductive interference), empirical tests of this prediction remain scarce. Here, we combined experimental and theoretical approaches to investigate the dynamics of two congeneric annual plant species (*Commelina*) experiencing both ecological competition and reproductive interference. Our empirically parameterized model revealed a large niche difference and mutual reproductive interference. However, the reproductive interference was not strong enough to generate pronounced contingency of community dynamics on initial conditions. Instead, coexistence was the most likely outcome regardless of initial species composition. Parameter exploration indicated that the weak contingency arose because reproductive assurance strategies of *Commelina* species buffered the minority disadvantage caused by reproductive interference. These findings provide empirical evidence that reproductive interference does not necessarily destabilize coexistence and that it may be outweighed by ecological niche and fitness differences. Our study highlights the importance of integrating ecological and reproductive processes towards better prediction of community dynamics.

**Significant statement:** Species interactions can both promote and undermine coexistence. Ecological niche differences can promote coexistence by favoring rare species, whereas reproductive interference can favor more abundant species and destabilize coexistence. How these opposing forces jointly shape community dynamics remains poorly understood. We combined experiments and mathematical approaches to examine their simultaneous effects on the coexistence of two closely related annual plants. Despite mutual reproductive interference, our empirically calibrated model showed that coexistence was the most likely outcome irrespective of initial species abundance. Reproductive interference had limited effects, likely because selfing before interspecific pollination provided reproductive assurance. Our findings demonstrate that reproductive interference does not necessarily destabilize coexistence and highlight the importance of integrating ecological and reproductive processes to predict community dynamics.

## Introduction

Understanding how biological interactions drive community dynamics and maintain species diversity is a fundamental question in ecology (Chesson, 2000; Simha et al., 2022). Classical coexistence theory posits that interspecific niche differences—arising from ecological mechanisms such as resource and predator partitioning—are necessary for stable coexistence because they make intraspecific competition stronger than interspecific competition, thereby generating negative frequency dependence in population growth (Chesson, 2000; Adler et al., 2007). For species to coexist, moreover, these niche differences must be large enough to offset differences in their baseline competitive ability. Numerous empirical studies have accordingly demonstrated evidence of interspecific niche differences (Harpole & Suding, 2007; Shinohara & Ohsaki, 2025) and compared them with fitness differences to assess the likelihood of coexistence (Godoy & Levine, 2014; Kraft et al., 2015).

Species also interact with each other through non-ecological, reproductive interactions, defined as behaviors associated with mate acquisition, which is potentially a more decisive driver of community dynamics (Kyogoku, 2015; Gómez-Llano et al., 2021). This is primarily because the negative interactions arising from imperfect species recognition and resultant interspecific mating—known as reproductive interference—generate positive frequency dependence in population growth, such that the adverse effect is disproportionally stronger on rarer species interacting with more abundant one (Kuno, 1992; Kyogoku, 2015; Weber & Strauss, 2016). In plants, for example, interspecific pollen transfer reduces reproduction opportunities, with the loss being more pronounced in rarer species (Levin & Anderson, 1970; Mitchell et al., 2009; Moreira-Hernández & Muchhala, 2019). Accumulating evidence suggests that reproductive interference plays evident roles in natural ecosystems, driving species exclusion, biological invasion, and distributional patterns both in animals (Thum, 2007; Kishi et al., 2009) and plants (Takakura et al., 2009; Whitton et al., 2017; Christie & Strauss, 2020).

When ecological competition and reproductive interference operate simultaneously, species may experience both advantages and disadvantages of being rare: the former resulting from interspecific niche differences and the latter from reproductive interference. Theory predicts that the interplay of positive and negative frequency dependence often results in multiple stable states, with the realized state depending on initial conditions (Yoshimura & Clark 1994; Kishi & Nakazawa, 2013; Schreiber et al., 2019). Nonetheless, empirical investigations of these combined effects remain limited. This gap partly reflects a long-noted, yet still persistent isolation between empirical studies in the two fields (Gómez-Llano et al., 2021). Studies on species coexistence have primarily focused on ecological niche and fitness differences (e.g., Godoy & Levine, 2014; Kraft et al., 2015), and rarely explicitly consider processes affecting reproduction, with only a few exceptions (Lanuza et al., 2018; Johnson et al., 2022; Cruz et al., 2025). On the other hand, although studies of reproductive interference sometimes examine resource competition (Takakura et al., 2009; Runquist & Stanton, 2013; Christie & Strauss, 2020), they do not evaluate how these processes shape long-term community dynamics. Consequently, although their prevalence in nature and theoretical appreciation, the combined effects of ecological competition and reproductive interference on community dynamics and species coexistence remain poorly understood.

Negative impacts of reproductive interference may be mitigated by various mechanisms, which are found widely both in animals and plants (Gröning & Hochkirch, 2008; Moreira-Hernández & Muchhala, 2019). This can be achieved either through reducing interspecific reproductive interactions or through securing intraspecific mating even with the opportunities of interspecific mating (Morales & Traveset, 2008; Moreira-Hernández & Muchhala, 2019). In plants, a typical example for the former mechanism is interspecific segregation of flowering phenology (Waser, 1978; Martin & Willis, 2007), while the latter involves self-pollination (Fishmand & Wyatt, 1999; Brys et al., 2016). Both types of mechanisms can alleviate the cost of rarity due to reproductive interference; even when such mechanisms are incomplete—for example, when flowering period partially overlap (Waser, 1978)—it may still relax reproductive interference and mitigate the minority disadvantage (Moreira-Hernández & Muchhala, 2019). However, the extent to which multiple simultaneous mechanisms counteract reproductive interference is poorly quantified, particularly in terms of their effects on community dynamics.

In this study, we aimed to quantify the joint effects of ecological competition and reproductive interference on the coexistence of two congeneric annual plants, *Commelina communis* and *C. c.* f. *ciliata*. Both species inhabit roadsides and farmlands, where they form single- or mixed-species patches locally and coexist in an intermingled state at mesoscales. Because they share pollinators due to morphological similarity of flowers (Fig. 1a) and do not hybridize, these species experience mutual reproductive interference (Katsuhara & Ushimaru, 2019). Nonetheless, they may alleviate the negative effect of reproductive interference through prior selfing (Katsuhara & Ushimaru, 2019; Katsuhara et al., 2024), in which individual flowers are self-pollinated before bud opening, thereby securing seed production even when surrounded by heterospecific individuals. In this system, we predicted that (1) because of ecological similarity, their dynamics would be close to neutral in terms of ecological competition (i.e., niche and fitness differences are small); (2) they would experience reproductive interference and exhibit positive frequency dependence in seed production; and (3) this effect would be incomplete because of prior selfing that ensures seed production when rare. Therefore, we anticipated that (4) reproductive interference would not be sufficiently strong to generate contingency on initial frequencies, such that community dynamics largely follow outcomes predicted solely from ecological competition. Finally, (5) we quantified the extent to which prior selfing mediates contingency in community dynamics, in comparison with phenological segregation as another potential mechanism mitigating reproductive interference.

**Figure. 1.**
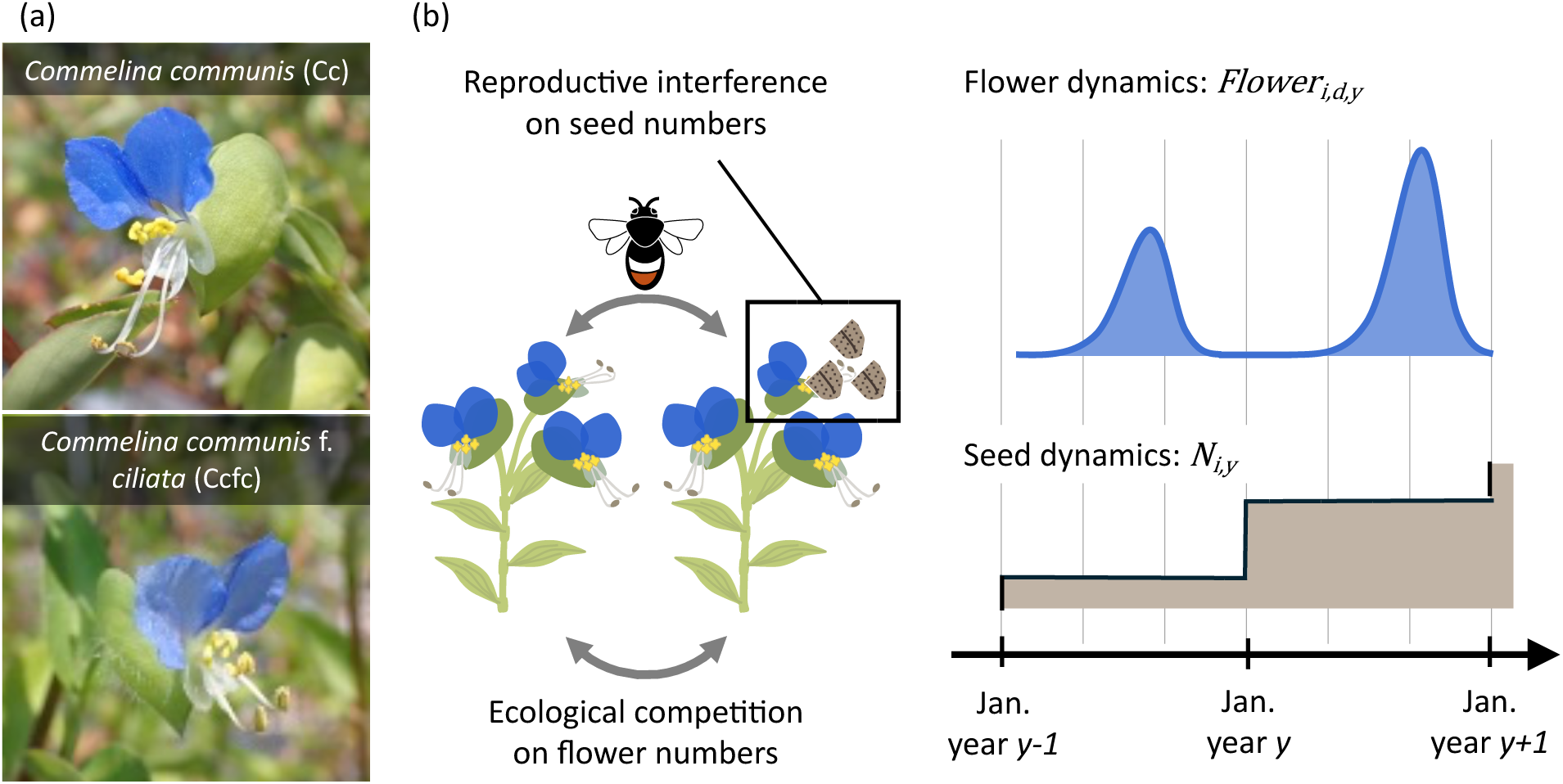
(a) Flowers of *Commelina communis* (Cc) and *C. c.* f. *ciliata* (Ccfc). The two species are morphologically very similar, with the most conspicuous difference being the presence or absence of cilia on bracts. (b) Schematic overview of our model. We assumed that ecological competition reduces the number of flowers produced by individuals over the growing season in a density-dependent manner, whereas reproductive interference affects the number of seeds produced per flower in a frequency-dependent manner. We explicitly modeled seasonal dynamics of flower production (*Floweri,d,y*) and interannual dynamics of seed abundance (*Ni,y*).

## Results

To examine the competitive dynamics of two *Commelina* species (*Commelina communis* [Cc] and *C. c.* f. *ciliata* [Ccfc]), we combined modeling and experimental approaches. We first developed a biologically realistic population dynamics model that explicitly incorporates ecological and reproductive processes of these species (Fig. 1b). We assumed that ecological competition reduces flower production in a density-dependent manner, whereas reproductive interference affects seed production per flower in a frequency-dependent manner (as in Runquist & Stanton, 2013; Christie & Strauss, 2020). The model considers seasonal dynamics of flower abundance, together with interannual dynamics of seed numbers, reflecting the fact that *Commelina* individuals produce one-day flowers continuously over several months each year.

We empirically parameterized the model through a series of field experiments. To quantify the density-dependent competition affecting flower numbers, we grew Cc and Ccfc in monocultures and mixtures across a wide range of densities in experimental pots (Table S1). We fitted a Ricker competition model to the relationship between flower production and plant density to estimate the competition parameters (Fig. S1). Our empirically calibrated model indicated that the coexistence of Cc and Ccfc was the most likely outcome when the number of seeds per flower was fixed for each species—a typical assumption in coexistence studies considering only ecological competition without reproductive interference. Across posterior draws of the parameter estimates, Cc and Ccfc were predicted to coexist in 89.5% of cases, whereas Cc and Ccfc were excluded in 6.9% and 3.4% of cases, respectively (Fig. 2). The high probability of coexistence reflects a sufficiently large niche difference arising from stronger intraspecific competition than interspecific competition (Fig. S2).

**Figure. 2.**
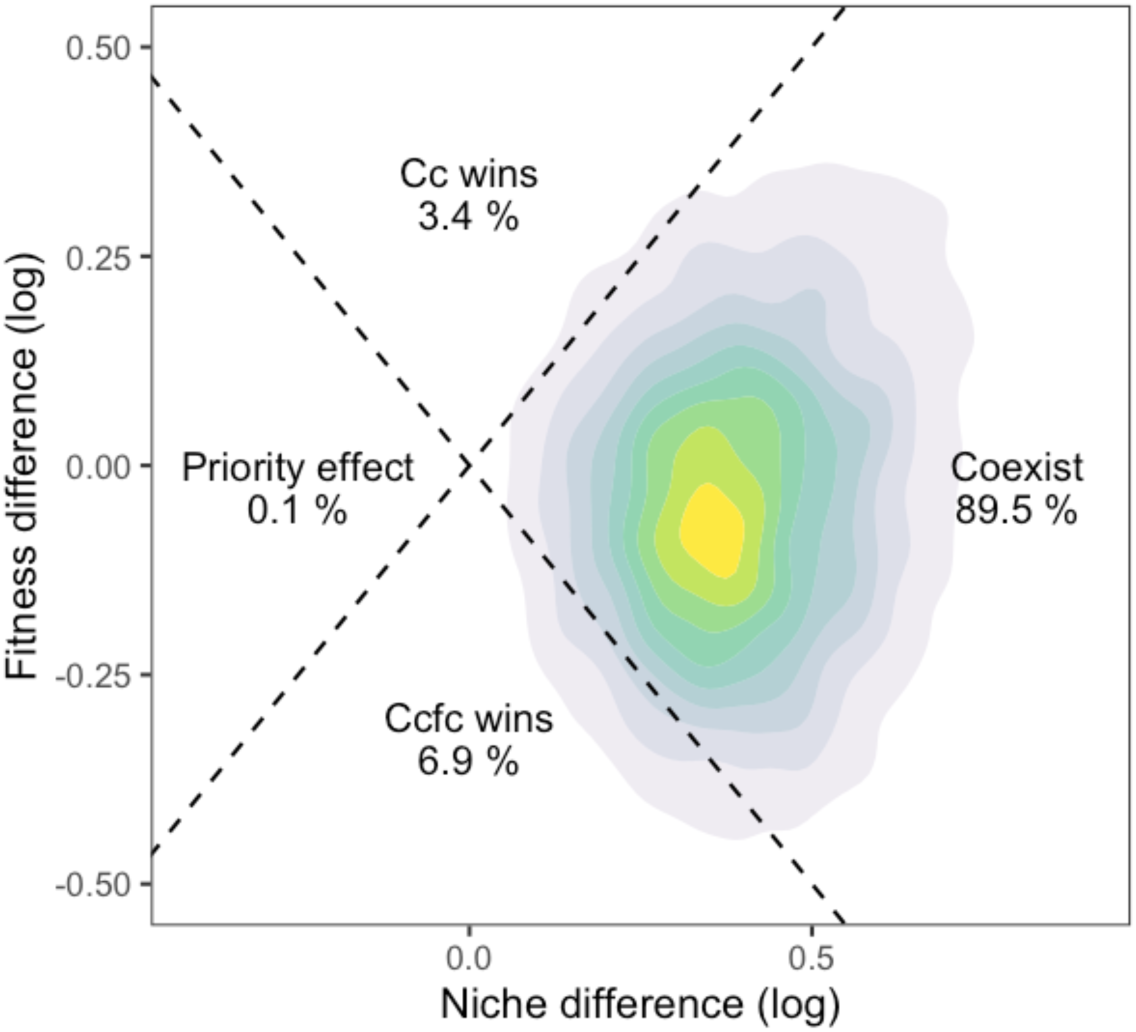
Predicted competitive outcomes of *Commelina communis* (Cc) and *C. c.* f. *ciliata* (Ccfc) based on the balance between niche and fitness differences. Broken lines indicate the theoretical boundaries separating competitive outcomes: coexistence, competitive exclusion of either Cc or Ccfc, and the priority effect. Shaded contours represent the posterior distribution of niche and fitness differences calculated from 1000 posterior draws of the model parameters. Percentages indicate the proportion of posterior draws falling within each outcome region.

We next estimated the strength of reproductive interference by examining the relationship between the number of seeds per flower and the frequency of conspecific flowers. We conducted a pollination experiment in which some of the experimental pots were enclosed within greenhouses, each containing a hive of bumblebees (*Bombus ignitus*). We also prepared a bumblebee-free greenhouse, in which selfing was the only means of pollination. During the pollination experiment period, different combinations of pots were introduced into the greenhouses each day to generate a full range of Cc and Ccfc flower frequencies within a greenhouse. We marked several flowers, let them be pollinated and examined their seed production a few weeks later. We found that, for both species, seed production per flower increased with the frequency of conspecific flowers within a greenhouse at the time of pollination (Fig. 3a), indicating mutual reproductive interference. The strength of reproductive interference was asymmetrical between these species, with the slope being larger for Cc (posterior mean: 0.89; 95% CI: [0.43, 1.33], Fig. 3b) than Ccfc (0.50; [0.03, 0.98]). Despite this frequency dependence, both species produced certain number of seeds even when conspecific frequency was close to zero (i.e., intercepts are 1.13 for Cc and 0.33 for Ccfc, Fig. 3a). This likely reflects reproductive assurance mechanisms that permit seed production without outcrossing, such as selfing. To examine this mechanism, we conducted an additional emasculation experiment in a bumblebee-free greenhouse where stamens were removed from flowers shortly after their opening so that prior selfing was the only possible means of pollination. Seed production resulting from prior selfing was greater in Cc (1.67 seeds produced per unpollinated flower) than in Ccfc (0.46 seeds) (Fig. S3), consistent with the larger intercept for Cc in Fig. 3a.

**Figure. 3.**
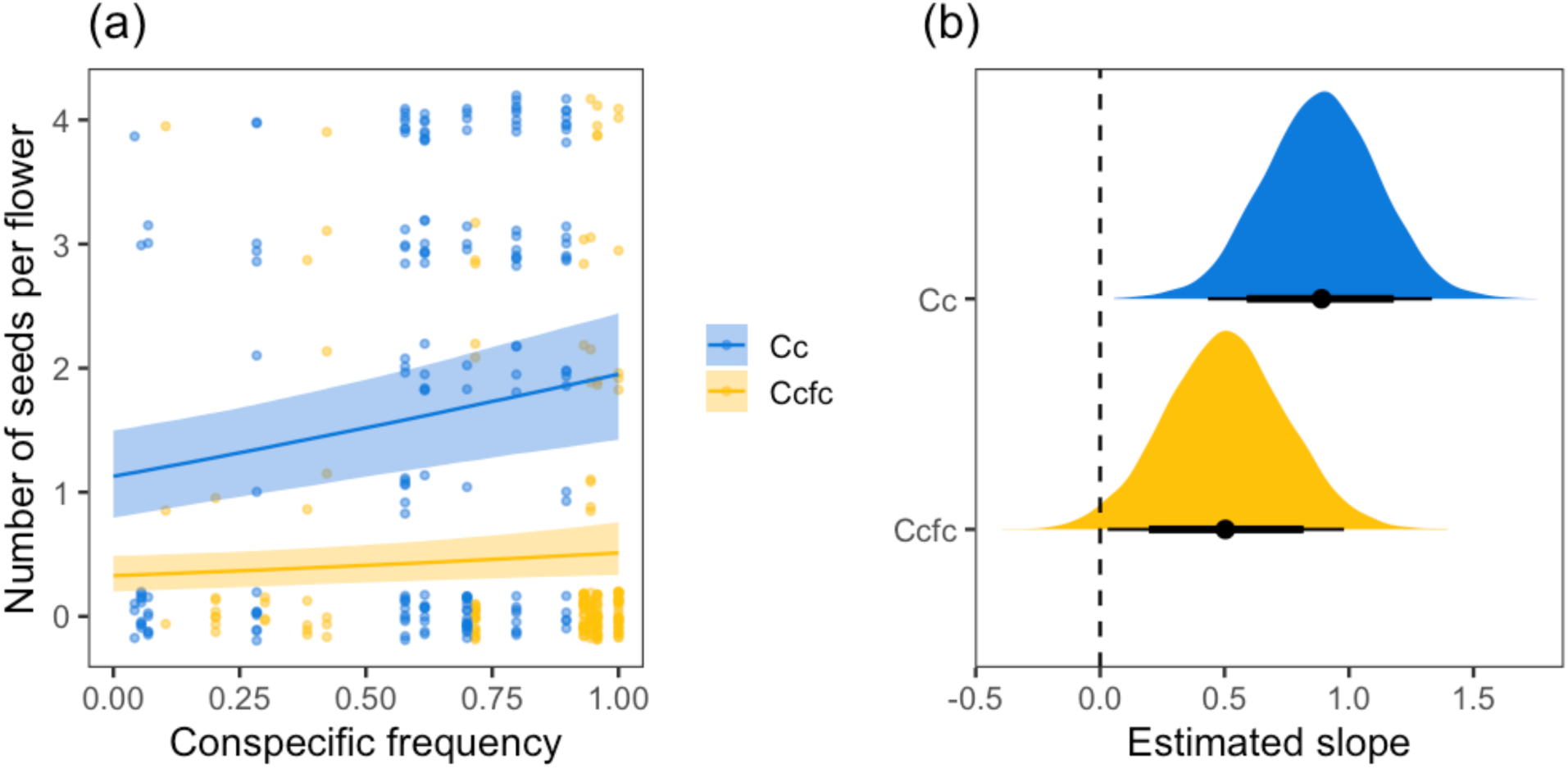
(a) Relationship between the number of seeds produced per flower and the frequency of conspecific flowers in the greenhouse at the time of pollination. Points represent individual flowers, and lines and shaded areas indicate posterior median predictions and 95% credible intervals from the regression models. (b) Posterior distributions of the estimated slopes for each species. Points indicate posterior medians, and thick and thin bars indicate80% and 95% credible intervals, respectively.

We then simulated the fully parameterized community dynamics model incorporating both ecological competition and reproductive interference. We ran simulations using 1000 parameter sets drawn from the posterior distributions to account for parameter uncertainty. We also considered three initial species frequencies—Cc dominant, equal frequency, and Ccfc dominant—to evaluate the dependence of community dynamics on initial conditions. We observed several qualitatively distinct dynamics. For some parameter sets, community dynamics initiated from the three frequencies converged to the same equilibrium state, either coexistence or exclusion of one species; we refer to these as *Global* outcomes (Fig. 4a). For other parameter sets, trajectories diverged depending on initial conditions, such that coexistence or exclusion of either species became possible depending on the initial frequency (*Contingent* outcomes, Fig. 4b). Among the 1000 parameter sets, global coexistence was the most common outcome (59.2%, Fig. 4b), followed by global exclusion of Ccfc by Cc (25.6%) and of Cc by Ccfc (5.1%). Contingent outcomes were less frequent, including coexistence or exclusion of Ccfc (8.4%), coexistence or exclusion of Cc (0.7%), and exclusion of either Cc or Ccfc (1.0%).

**Figure. 4.**
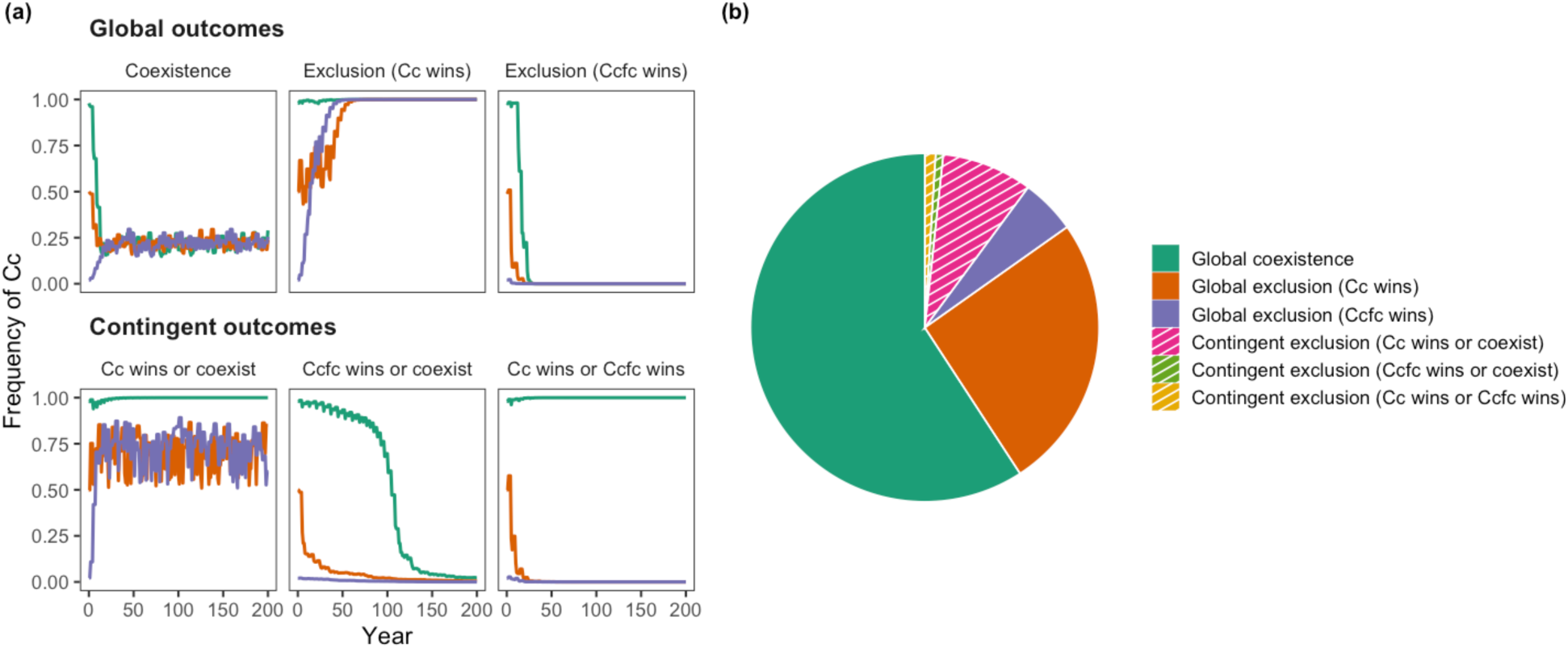
(a) Representative simulations of the interannual dynamics of the frequency of *Commelina communis* (Cc) within a community. Each panel corresponds to a different set of parameters drawn from the posterior distribution, and lines represent the simulated trajectories initiated from different frequencies. Different parameter sets produced qualitatively distinct dynamics, some independent of initial conditions (*global* outcomes: coexistence and exclusion) and others contingent on them (*contingent* outcomes). (b) Proportions of the dynamical outcomes exemplified in (a) across 1000 posterior parameter sets. Contingent dynamics are indicated by diagonal hatching.

We further quantitatively assessed the extent to which reproductive interference is mitigated by two potential mechanisms: phenological segregation, which reduces interspecific reproductive interactions, and selfing, which allows individual flowers to produce seeds even when opportunities for interspecific mating are abundant. First, we manipulated the parameter values of flowering peaks to consider scenarios where the flowering phenologies of two species either strongly segregated or substantially overlapped (SI Appendix Supporting Methods). Second, we systematically varied the “selfing rates” (i.e., the intercepts in Fig. 3a) from near zero (0.001) to the maximum seed production of each species (SI Appendix Supporting Methods). These two extremes corresponded to nearly complete reproductive interference, in which individuals are unable to produce seeds when rare, and the absence of reproductive interference, in which conspecific frequency does not affect seed production, respectively. Using these modified parameters, we simulated community dynamics starting from three different conditions and assigned the resulting dynamics to either of the *global* or *contingent* outcome categories. Contingent dynamics became more likely as selfing rates decreased and phenological overlap increased (Fig. 5), consistent with stronger reproductive interference under these conditions. The result further suggested that the observed selfing rates were sufficiently high that contingent dynamics remained unlikely even with substantial phenological overlap, highlighting the critical role of selfing in mitigating the effect of reproductive interference.

**Figure. 5.**
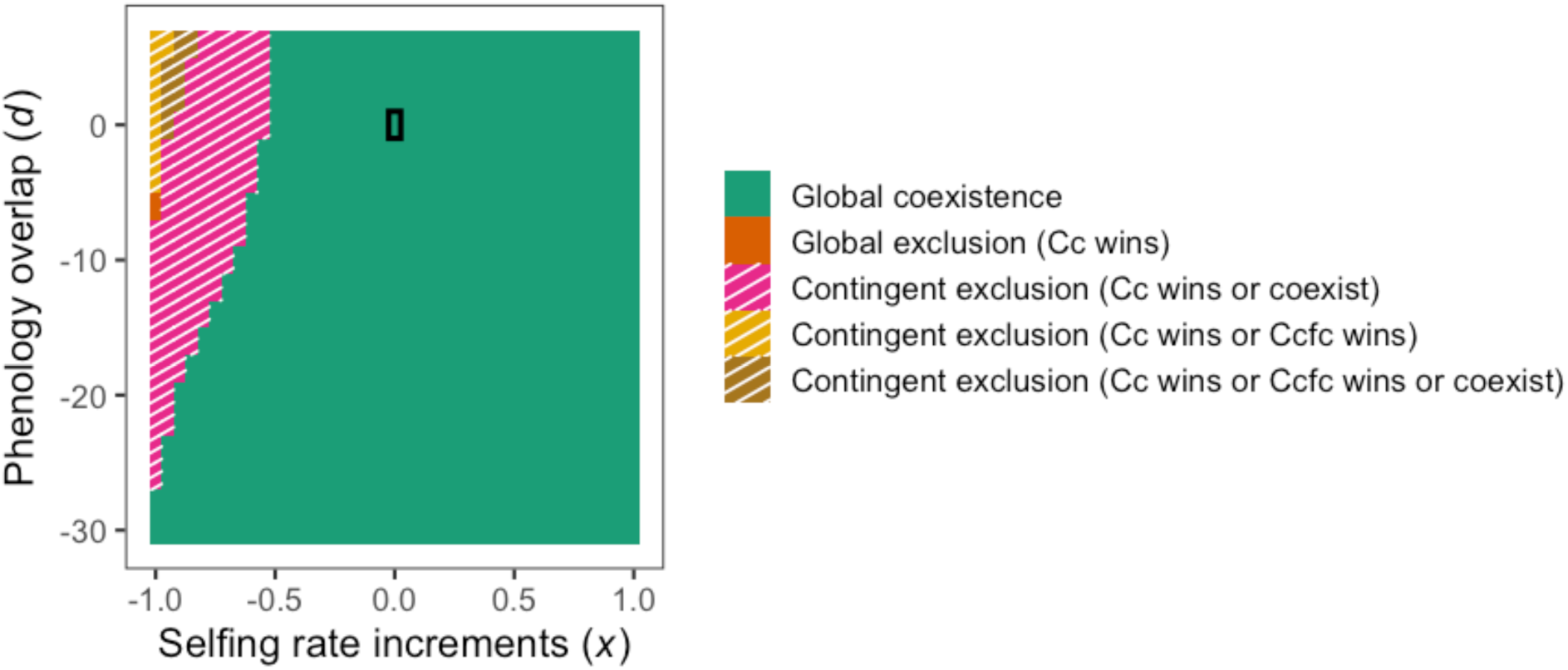
Predicted dynamical outcomes under varying selfing rates (the regression intercept in Fig. 3a) and degrees of interspecific phenological segregation (see SI Appendix Supporting Methods for details). The selfing rate increment parameter (*x*) controlled the strength of reproductive interference: reproductive interference was absent when *x* = 1, whereas it was nearly complete (i.e., each species produced no seeds when rare) when *x* = –1. Interspecific phenological overlap was greatest when *d* = 7 and smallest when *d* = –30. *x* = 0 and *d* = 0 correspond to the empirically estimated values (black rectangle). Parameter combinations resulting in contingent dynamics are indicated by diagonal hatching.

## Discussion

Theory suggests that ecological competition and reproductive interference jointly shape complex community dynamics arising from the combination of negative and positive frequency dependence in population growth. Notably, their simultaneous operation is predicted to generate contingent dynamics, with multiple ecological states—exclusion of either competitor or coexistence of multiple species—being determined by initial conditions (Yoshimura & Clark 1994; Kishi & Nakazawa, 2013; Schreiber et al., 2019). Nonetheless, reflecting the historically separate development of these two fields, empirical investigation of their joint effects has been surprisingly scarce (Gómez-Llano et al., 2021). The present study fills this gap by demonstrating that, despite the evidence of mutual reproductive interference between two congeneric annual plant species (Fig. 3), reproductive interference had little appreciable impact on community dynamics: The dynamics projected by our empirically calibrated model largely followed the outcome predicted from ecological competition alone (i.e., coexistence, Fig. 2), regardless of initial frequency (Fig. 4). This result suggests that the destabilizing effect of reproductive interference may be outweighed by ecological niche and fitness differences, highlighting the necessity of integrating ecological and reproductive interactions to better understand species coexistence in nature.

Reproductive interference occurs in a frequency-dependent manner (Levin & Anderson, 1970; Kuno, 1992) and is therefore often considered to be more influential in driving ecological dynamics than competition, which acts in a density-dependent manner (Kishi et al., 2009; Runquist & Stanton, 2013; Christie & Strauss, 2020). Indeed, research on reproductive interference has linked the observation of positive frequency dependence in fitness measures to ecological patterns such as species exclusion, biological invasion, and species distributions (Takakura et al., 2009; Runquist & Stanton, 2013; Christie & Strauss, 2020). Our findings call for a more nuanced view of this conventional interpretation. Like these previous studies, we detected positive frequency dependence in seed production between two congeners, a hallmark of reproductive interference (Fig. 3). However, our results indicated that the effects of reproductive interference can be outweighed by ecological competition, to the point that contingent dynamics predicted from reproductive interference were rarely observed. This suggests that demonstrating the presence of reproductive interference *per se* is insufficient to conclude that it drives ecological patterns in the field; rather, comparing the relative contributions of reproductive interference and ecological competition is necessary. Although several studies have conducted such comparison in certain ways (e.g., Kishi et al., 2009; Runquist & Stanton, 2013; Christie & Strauss, 2020), scenario analyses of long-term community dynamics, as we conducted here (e.g., with and without reproductive interference: Fig. 2 versus Fig. 4) can provide particularly informative evidence.

A key finding of this study is that the two *Commelina* species could produce seeds even when rare (i.e., intercepts were greater than zero in Fig. 3). This explains why the projected community dynamics were largely insensitive to initial frequency (Fig. 4): because the minority cost imposed by reproductive interference was relatively small, it was offset by the advantage of being rare conferred by the niche difference. Consistent with this interpretation, reducing the intercept values shifted the projected dynamics from global outcomes to contingent ones (Fig. 5). We referred to this intercept as the selfing rate in the Results section simply because higher selfing rates should result in greater seed production in the absence of conspecific pollination. Nonetheless, the intercept may also reflect contributions from other mechanisms that lead to similar biological outcomes. For example, pollinator flower constancy, where pollinators tend to visit flowers of the same species disproportionately often relative to its availability, could increase the likelihood of intraspecific pollination even when only a few conspecific individuals are present (Levin & Anderson, 1970; Moreira-Hernández & Muchhala, 2019). Also, even when pollinators move between flowers of different species, intraspecific pollination can be ensured if pollen adheres to different parts of pollinator’s body (Huang & Shi, 2013). This mechanism, however, is unlikely to be important in our *Commelina* system given the similarity in thier floral mechanics (e.g., positions of the anther and stigma). Disentangling the contributions of these mechanisms would be difficult. Nonetheless, the observed intercepts were of a similar order of magnitude to the prior selfing rates independently measured in this study (Fig. S3), supporting our interpretation that the intercept provides a reasonable proxy for selfing rate. This interpretation is also consistent with a previous study reporting higher selfing rates in Cc than in Ccfc (Katsuhara & Ushimaru, 2019).

Species often possess multiple mechanisms that can potentially alleviate the negative impact of reproductive interference (Morales & Traveset, 2008; Moreira-Hernández & Muchhala, 2019). Although individual mechanisms, such as phenological or mechanical segregation, have been shown to be effective (Waser, 1978; Huang & Shi, 2013), the presence of a mechanism does not necessarily imply it is the most important when multiple mechanisms operate. Assessing the relative effectiveness of such simultaneous mechanisms therefore remains a challenge. One approach is to perform interspecific comparisons. For example, Christine & Strauss (2018) compared the strength of several reproductive barriers and genetic distance across 39 congeneric species pairs, suggesting that prezygotic barriers, such as phenological segregation, were more pervasive early in speciation, whereas postzygotic barriers became stronger with evolutionary time. Our approach provides an alternative framework, with a greater focus on ecological outcomes. We demonstrated that selfing would more easily reduce the contingency of community dynamics than flowering phenological segregation (Fig. 5). Interestingly, this finding is consistent with Christine & Strauss (2018), suggesting that phenological segregation may become less effective once postzygotic reproductive barriers have been established. Our approach therefore provides a complementary perspective for understanding the ecological consequences of reproductive strategies.

The presence of a sufficiently large niche difference between Cc and Ccfc was unexpected (Fig. 2), given their similarity in habitat preference (Fig. S2 in Katsuhara & Ushimaru, 2019) and morphology–the only visually appreciable difference is the presence (Cc) or absence (Ccfc) of cilia on their bracts enclosing inflorescences (Fig. 1a), whose ecological function is unclear. Nonetheless, we detected interspecific differences in several ecological traits, which collectively indicate that Ccfc is a resource-acquisitive strategist, with higher specific leaf area, leaf nitrogen content, photosynthetic efficiency, and specific root length, whereas Cc is more conservative, with higher leaf dry mass content and leaf C:N ratio (Fig. S4). We also found that Cc has greater water-use efficiency, as indicated by higher δ13C value (Fig. S4). Consistent with this finding, a recent study found that Cc has lower stomatal density (Shimonai et al., 2024), which may confer an advantage in water-use efficiency by reducing water loss through leaf transpiration. Therefore, these differences in resource-use strategies may underpin the niche difference between Cc and Ccfc, despite not being readily apparent from external morphology. Indeed, substantial niche differences have been documented between many congeneric pairs (e.g., Ocampo-Ariza et al., 2018; Blackford et al., 2020), underscoring that phylogenetic proximity does not necessarily imply ecological similarity.

The primary aim of this study was to facilitate the empirical integration of ecological and reproductive processes. To this end, we developed a biologically realistic community dynamics model, which allowed us to isolate these two processes and compare their relevance to long-term dynamics. Nonetheless, we made several simplifying assumptions that should be considered when extrapolating our findings to real-world systems. First, our model assumed that pollinator abundance was always sufficient to induce reproductive interference. However, pollinator abundances are not constant either spatially or temporally in natural ecosystems (Lanuza et al., 2018), indicating that the strength of reproductive interference could also vary dynamically. Including pollinator dynamics into the model (e.g., Bartomeus et al., 2021) would expand the scope, for example, to projecting the consequences of future pollinator loss (Johnson et al., 2022). Second, we assumed that seed production per flower was solely influenced by reproductive interference, not by competition. We adopted this assumption based on *Commelina* species’ biology that each flower has four ovules, a number strongly regulated during flower development and is therefore unlikely to vary in response to competition. Nonetheless, it is possible that realized seed production is affected by resource availability. Indeed, the maximum seed production observed in this study (approximately 1–2 seeds per flower, Fig. 3) was lower than that reported in field conditions (2–3 seeds; Katsuhara & Ushimaru, 2019). This discrepancy can be explained by the difference in the resource availability. *Commelina* species commonly occur along agricultural margins where nutrient availability is relatively high, whereas in this study we added nutrients only at the beginning of the experiment. Thus, in our experiment, flowers may have failed to produce seeds even when successfully pollinated because of the lack of nutrients, particularly phosphorus. This possibility suggests that competition for resources and reproductive processes may interact, a synergetic effect that warrants further investigation (Cruz et al., 2025). Finally, our model considers only local dynamics, despite that these species are typically distributed patchily within a landscape, forming a metacommunity (Katsuhara and Ushimaru, 2019; Fujiwara et al., 2025). Their coexistence and natural distributions are therefore determined by the interplay between local biological interactions and regional processes such as dispersal and environmental filtering processes (Shinohara and Katsuhara, 2025). However, disentangling their collective roles remains a major challenge in the study of species coexistence in general (Ellner et al., 2024).

## Material and Methods

### Study system and focal species

We focused on two congeneric annual plants *Commelina communis* (Cc) and *C. c.* f. *ciliata* (Ccfc). They possess similar ecological characteristics including habitats (both are found around agricultural areas and along roadsides), morphology (similar in plant size and shapes of leaves and flowers), and flowering phenology (both peak in summer, Fig. S5). During growing seasons, individuals produce several one-day flowers every day, which open after sunrise and close by afternoon. Successfully pollinated flowers produce seeds up to four, the upper limit being determined by the number of ovules per flower. Importantly, these species are self-compatible; self-pollination occurs when the stamens and pistil within the same flower contact each other before flower opening (i.e., prior selfing) or during flower closure (delayed selfing). It is also noteworthy that both species are andromonoecious, meaning that individuals bear both perfect flowers with stamens and a pistil and staminate flowers with only stamens. Only perfect flowers contribute to seed production, whereas both flower types can be pollen donors.

Because of the floral similarity, they share almost identical pollinator niches, for which interspecific pollination is observed very often (Katsuhara and Ushimaru, 2019). Interspecific outcrossing substantially reduces their seed production (Katsuhara and Ushimaru, 2019), the incompatibility being attributable to different chromosomes numbers (Cc: 2n = 88; Ccfc: 2n = 46, Shimonai et al., 2024). Consequently, reproductive interference occurs between them, such that their seed production positively correlates with conspecific flower frequency (Katsuhara and Ushimaru, 2019).

### Population dynamics model

We developed a population dynamics model specifically designed for this *Commelina* system that describes both interannual dynamics of seed abundance (*Ni,y*) and seasonal dynamics of flower abundance (*Floweri,d,y*) (Fig. 1). The model consists of two components. We assumed that ecological competition reduces the total number of flowers produced per individual over the growing season, while reproductive interference affects the number of seeds produced per flower (as in Runquist and Stanton, 2013; Christie and Strauss, 2020). Model details are provided in SI Appendix Supporting Methods and briefly summarized in Table S2.

### Competition experiment

To parameterize our competition model, we planted germinated individuals into experimental pots with varying densities (Table S1), including single-individual, monoculture, and mixed-culture treatments. Details of seed origin and planting and growing conditions are provided in SI Appendix Supporting Methods. To capture their flowering phenology and lifetime flower production, we counted the number of flowers of each species in each pot as many days as possible throughout the growing season (during June–November 2024).

### Pollination experiment

For parameterization of the reproductive interference model, we conducted a pollination experiment using three greenhouses (4.5m × 2.2m × 2.1m height), two of which each contained a hive of bumblebees (*Bombus ignitus*), while the third contained none. The bumblebee hives were purchased from a local agricultural supplier.

The experimental procedures were as follows. Each evening during the period of pollination experiment (from September to October), we placed five experimental pots into each greenhouse. The combination of the pots was varied arbitrarily each day to generate the full range of frequencies of Cc, from 0% (only Ccfc flowers present) to 100% (only Cc) (Fig. 3). On the following morning, we counted the total number of perfect and staminate flowers in each greenhouse, selected up to five Cc or Ccfc perfect flowers from each pot, and marked them with small plastic tape tags. We then opened the bumblebee hives for 2 hours, allowing bees to transport pollen from flower to flower. A few weeks later, we assessed the seed production of marked flower after maturation. In the greenhouse without a bumblebee hive, the same procedures were conducted except for the release of bumblebees.

During the period when bumblebees were released, we recorded their visits to flowers (either Cc or Ccfc) in each greenhouse for 30 minutes. After excluding observations with insufficiently low visitation frequency (which were omitted from the subsequent analyses), the mean number of flower visits was 317 times in 30 minutes. Dividing the visit number by the flower number within a greenhouse, each flower received an average of 8.1 visits during the 2-hour period when the hives were open, which appeared sufficient for pollen transfer. Indeed, we occasionally confirmed that no pollen remained on stamens after bee release. We additionally conducted an emasculation experiment where stamens were removed after flower opening and flowers were left unpollinated in the bumblebee-free greenhouse. This allows us to examine the extent of prior selfing.

### Parameter estimation

Parameters of our population dynamics models were estimated from the experiments. Details of the statistical models are provided in SI Appendix Supporting Methods. In short, we used several linear or nonlinear Bayesian regression models and obtained the posterior draws of the parameters. For the Bayesian inference, we used the *brm* function in the *brms* package (Bürkner, 2017) in R (R Core Team 2024).

### Community dynamics simulations

With the empirically parameterized model, we simulated the dynamics of Cc and Ccfc under three scenarios. First, we considered a scenario where ecological competition was the sole driver of community dynamics. This was implemented by fixing the number of seeds produced per flower at its maximum value predicted by our regression model (i.e., the value when conspecific frequency is one in Fig. 3: 1.99 for Cc and 0.521 for Ccfc). This approach mirrors the typical practice in coexistence studies that consider only ecological competition and ignore reproductive interference. Under this scenario, competitive outcomes can be analytically determined by comparing the niche and fitness differences between species (SI Appendix Supporting Methods). To account for uncertainty in parameter estimation, we calculated the niche and fitness differences from 1000 parameter sets drawn from the posterior distributions.

Second, we numerically simulated the community dynamics model including both ecological competition and reproductive interference for 100 years. To evaluate the potential contingency of community dynamics on initial species frequencies due to positive frequency dependence, simulations were initiated from three combinations of seed abundances of Cc and Ccfc: 49:1; 25:25: and 1:49. For each simulation run, we classified the competitive outcome (coexistence or exclusion of either Cc or Ccfc) according to the frequency in the final year. Again, we used 1000 parameter sets for the simulation and classified each parameter set as yielding either *global* or *contingent* dynamics (see Fig. 4) based on the correspondence among competitive outcomes across the three initial conditions. Further details of the simulations and outcome classification are provided in SI Appendix Supporting Methods.

Finally, we manipulated parameter values so that we can systematically vary the difference in peak flowering phenology between Cc and Ccfc, as well as the intercepts of the regression between seed production and conspecific flower frequency (Fig. 3a). These manipulations correspond to hypothetical changes in the degree of interspecific phenological segregation and selfing rates. For each modified parameter combination, we again simulated community dynamics with three initial conditions and assigned the dynamical outcome (*global* or *contingent* one) based on the correspondence among their competitive outcomes (see SI Appendix Supporting Methods for details).

### Trait measurement

We measured functional traits of Cc and Ccfc individuals grown separately from the competition and pollination experiments. We sowed seeds harvested from the competition experiment in the following year of the experiment into plastic pots, and measured several functional traits from 13 Cc and 10 Ccfc individuals. We selected traits associated with plant resource acquisition strategies: leaf dry mass content, specific leaf area, photosynthetic efficiency (SPAD), leaf nitrogen content, leaf C:N ratio, and specific root length. We also measured leaf δ^13^C and δ^15^N, which are commonly used as indicators of water-use efficiency and nitrogen-use strategy, respectively. All measurements followed standard protocols (Perez-Harguindeguy et al., 2016). Details are provided in SI Appendix Supporting Methods.

## Acknowledgement

The authors thank Kentaro Takagi and Kazuharu Ohashi for the advice on rearing bumblebees, Fujio Hyodo and Yuko Miyazaki for the help in the trait measurements, and Teruki Inoue, Hiyori Sasaki, Takeo Doi, Takuto Kaneko, Akira Matsumoto, and Rie Yoshinami for the assistance in the preparation of the experiments. This work was supported by Grant-in-Aid for JSPS Fellows 23KJ0077 (to NS), Joint Usage/Research Grant of Center for Ecological Research (2024jurc-cer03), Kyoto University, Inamori Research Grants, and JSPS KAKENHI Grant Number 21K17914 (to KRK).

