## Supplementary material for "Coexistence of annual plants under competition and reproductive interference": SI Appendix

**Authors**: Naoto Shinohara[1†], Akira Yamawo[2], Koki R. Katsuhara[3†]
[1]Okinawa Institute of Science and Technology, [2]Kyoto University, [3]Okayama University
†: These authors contributed equally

**Contents**

**Supplementary methods**


1. Population dynamics model
2. Competition experiment
3. Trait measurement
4. Parameter estimation
    - 4.1. Estimation of population dynamics parameters
    - 4.2. Prior sensitivity of candidate models
    - 4.3. Seed germination and survival probabilities
5. Community dynamics simulations
    - 5.1. Derivation of the coexistence condition without reproductive interference
    - 5.2. Classification and examination of simulated dynamics
    - 5.3. Scenarios of varying degrees of selfing rates and phenology partitioning


**Supplementary materials**


Table. S1–S3, Figure. S1–S5

### Supplementary methods

#### *1. Population dynamics model*

We developed a population dynamics model that describes the seasonal dynamics of flower production and the interannual dynamics of seed abundance of two congeneric annual plants, *Commelina communis* (Cc) and *C. c.* f. *ciliata* (Ccfc). Reflecting their biology, our model assumed that ecological competition reduces the number of flowers produced by individuals, whereas reproductive interference influences the number of seeds produced per flower. See Table S2 for the brief summary of the model.

The interannual seed dynamics of species $i$ ($N_{i,y}$, $y$ denotes year) are described as:

$$N_{i,y+1} = (1 - g_i)s_i N_{i,y} + g_i N_{i,y} \sum_{d=d_{min}}^{d_{max}} p_i Flower_{i,d,y} Seed_{i,d,y}\,, \qquad (S1)$$

where $g_i$ and $s_i$ respectively represent the germination rate and the survival rate of ungerminated seeds of species $i$. $Flower_{i,d,y}$ is the total number of flowers—including perfect and staminate—produced by an individual of species $i$ on day $d$, and $Seed_{i,d,y}$ is the number of seeds produced by a perfect flower of species $i$ blooming on day $d$. $d_{min}$ and $d_{max}$ denote the beginning and end date of the flowering season, which were fixed at 1 June and 1 November, respectively, for both species. In the summation, $Flower_{i,d,y}$ is multiplied by $p_i$, the proportion of perfect flowers among all flowers (perfect and staminate) produced by species $i$, reflecting that only perfect flowers produce seeds. We confirmed that $p_i$ differed between the two species but was not substantially affected by other factors such as competition intensity or seed sources (Table S3); therefore, $p_i$ was fixed at 0.882 for Cc and 0.769 for Ccfc throughout the analyses. Consequently, the summation term in Eq. S1 represents the lifetime seed production of a germinated individual of species $i$ in year $y$.

Seasonal flower production was modelled in two steps. First, we assumed that competition with conspecific and heterospecific neighbors reduces $TotalFlower_{i,y}$, the total number of flowers (including perfect and staminate) produced by an individual of species $i$ over the growing season in year $y$. We assumed that the competitive effects follow the Ricker function, for the reason described later (Section *4.2. Prior sensitivity of candidate models*):

$$TotalFlower_{i,y} = \lambda_i \exp\left(-\alpha_{ii} g_i N_{i,y} - \alpha_{ij} g_j N_{j,y}\right), \qquad (S2)$$

where $\lambda_i$ represents the maximum flower production by an individual of species $i$, and $\alpha_{ii}$ and $\alpha_{ij}$

represent strengths of competitive effects from germinated conspecific (species *i*) and heterospecific (*j*) individuals, respectively. Then, $TotalFlower_{i,y}$, was distributed across flowering dates within the growing season according to a Gaussian probability function, yielding $Flower_{i,d,y}$, flower production by an individual of species *i* on day *d* in year *y*:

$$Flower_{i,d,y} = TotalFlower_{i,y} \frac{1}{\sqrt{2\pi\sigma_i^2}} \exp\left(-\frac{(d-m_i)^2}{2\sigma_i^2}\right), \qquad (S3)$$

with $m_i$ representing the peak flowering date and $\sigma_i$ the standard deviation of the gaussian flowering phenology curve of species *i*.

The number of seeds produced by a flower, $Seed_{i,d,y}$, was modelled as a function of the frequency of conspecific flowers ($f_{i,d,y}$) at the time of pollination, reflecting the effect of reproductive interference. Because per-flower seed production of these *Commelina* species is bounded between 0 and 4 (the number of ovules per flower), we adopted a sigmoid function:

$$Seed_{i,d,y} = 4\frac{\exp\left(a_i + b_i f_{i,d,y}\right)}{1+\exp\left(a_i + b_i f_{i,d,y}\right)}, \qquad (S4)$$

where

$$f_{i,d,y} = \frac{g_i N_{i,y} Flower_{i,d,y}}{g_i N_{i,y} Flower_{i,d,y} + g_j N_{j,y} Flower_{j,d,y}}. \qquad (S5)$$

In Eq. S4, $a_i$ determines the baseline of seed production when the conspecific frequency is zero, whereas $b_i$ represents the effect of increasing conspecific flower frequency. Therefore, $b_i$ is expected to be positive if reproductive interference occurs. Note that this formulation of reproductive interference differs from those conventionally used in theoretical studies, such as $f_{i,d,y} / (f_{i,d,y} + b_i f_{i,d,y})$ (e.g., Kuno, 1992, Schreiber et al., 2019). Our choice of Eq. S4 was motivated by the biology of *Commelina* species: individuals may produce seeds even when conspecific flowers are rare ($f_{i,d,y} \approx 0$) through reproductive insurance mechanisms, such as selfing. This possibility is not accounted for in the conventional formulations.

*2. Competition experiment*

Seeds of Cc and Ccfc were collected from mature individuals growing at edges of rice fields in the western part of Japan mainland (Okayama prefecture) in 2023 and stored at room temperature until the experiment. On 23 May 2024, after germination on seedling trays, seedlings were transplanted into 21 experimental pots (43 cm × 28.6 cm) filled with fertile soil (10 cm depth; N:P:K = 320:210:300 [mg/L]) at varying densities (Table S1): single-individual pots, monocultures containing multiple individuals (up to five individuals), and mixed cultures with multiple individuals (up to five in total). To increase replication, seedlings that emerged later were transplanted into additional eight experimental pots on 13 June 2024. For the same purpose, we also collected geminated individuals from field populations (Cc: the Center for Ecological Research, Kyoto University; Ccfc: edges of rice fields in Osaka Prefecture) and transplanted them into additional 20 experimental pots on 9 May 2024. In total, we had 49 experimental pots with multiple combinations of densities (Table S1), with seedlings originating from three source/planting date categories: early (field-collected seedlings transplanted on 9 May 2024), middle (seedlings germinated from collected seeds and planted on 23 May) and late (seedlings germinated from collected seeds and planted on 13 June) (Table S1). The effects of seedling source and planting date were accounted for in the following analyses.

We placed the experimental pots in a research field in the Center for Ecological Research of Kyoto University, Japan. The pots were watered regularly throughout the experiment period. These species produce multiple one-day flowers every day during the flowering season from June to October. To capture flowering phenology and estimate the total flower production, we counted the number of all flowers (including perfect and staminate) of each species in each experiment pot as frequently as possible (60 days during their flowering period). Note that the flower numbers were counted at *pot* level, rather than *individual* level, because these species form dense vegetation, making it infeasible to distinguish individuals within pots. To estimate the proportion of perfect flowers among both perfect and staminate flowers ($p_i$ in our model: Eq. S1), we separately counted the numbers of perfect and staminate flowers for each experimental pot occasionally (seven days during the period).

### *3. Trait measurement*

We measured functional traits of healthy individuals of Cc and Ccfc grown independently from the competition and pollination experiments. In 2025, seeds collected from individuals used in the competition experiment in 2024 were sown in seedling pots. Emergent seedlings were transplanted individually into 5-L pots filled with fertile soil (N:P:K = 320:210:300 [mg/L]), and 13 Cc and 10 Ccfc individuals were subsequently used for trait measurements. The pots were situated at the Okayama University experimental field. Following the standard protocols (Perez-Harguindeguy et al., 2016), we selected eight traits associated with leaf economics and water-use efficiency: photosynthetic efficiency (SPAD), specific leaf area (SLA), leaf dry mass content (LDMC), specific root length (SRL), leaf nitrogen content, leaf $\delta^{15}N$, leaf $\delta^{13}C$, and leaf C:N ratio. At peak growth (2 October), SPAD values were calculated as the average of five measurements taken from each of three randomly selected leaves per individual using a SPAD-502 chlorophyll meter (Konica Minolta, Tokyo, Japan). Another 10 leaves were sampled, kept moist and cool, and used to measure fresh mass and leaf area with a scanner (CANON CanoScan LiDE 400) on the same day. After drying at 70°C for 72 h, their dry mass was measured, and the leaves were pulverized for isotope analysis. Leaf nitrogen and carbon content, as well as leaf $\delta^{15}N$ and $\delta^{13}C$, were measured using an isotope ratio mass spectrometer (Delta V Advantage, Thermo Fisher Scientific Inc., Waltham, MA, USA) connected to an elemental analyzer following the method of Ogawa et al. (2010). At the end of growth (17 November), a portion of the root of each individual was carefully washed under running water, scanned using a scanner, and measured for length using ImageJ (Schneider et al., 2012). Root samples were then dried at 70°C for 72 h, and then their dry mass was measured.

### *4. Parameter estimation*

#### *4.1 Estimation of population dynamics parameters*

Eq. S2, S3, and S4 were empirically parameterized based on the competition and pollination experiments. Firstly, the flowering phenology parameters in Eq. S3 were estimated by fitting the observed dynamics of the number of all flowers for each experimental pot ($p$) and each species ($i$) to the equation. We used a nonlinear regression:

$$\text{Observed per-pot flower number}_{i,p,d} \sim Pois\left(fpeak_{i,p} \exp\left(-\frac{(d-m_{i,p})^2}{2\sigma_{i,p}{}^2}\right)\right), \qquad (S6)$$

with $fpeak_{i,p}$ representing the total number of flowers of species $i$ in pot $p$ produced at the phenology peak, and $m_{i,p}$ and $\sigma_{i.p}$ corresponding to the parameters in Eq. S3 but estimated separately for each pot. The day $d$ was converted to a numeric value centered to the median date of the observation ($d$ = 0 on August 27, 2024). We opted weakly informative priors for these parameters based on our knowledge on the phenology: $fpeak_{i,p}$ ~ Normal [40, 20] with restricted to be positive, reflecting that an individual produces around 40 flowers per day in maximum (Katsuhara & Ushimaru, 2019); $m_{i,p}$ ~ Normal [0, 30], as the peak was observed in the middle of the season; and $\sigma_{i.p}{}^2$ ~ Normal [900, 450] with only positive values allowed, to make the expected range of 95% probability interval of the flowering phenology approximately [–60, 60] (zero ± 2$\sigma_{i.p}$). We used the *brm* function in the *brms* package (Bürkner, 2017) in R (R Core Team 2024) and calculated the posterior distributions of these parameters from the Markov chain Monte Carlo sampling method with four chains and 4000 iterations, with 2000 used as a warm-up. We averaged $m_{i,p}$ and $\sigma_{i.p}$ values across estimates from all experimental pots to obtain species-specific $m_i$ and $\sigma_i$ values in Eq. S3, which was subsequently used for community dynamics simulation. We fixed these phenology parameters per species, ignoring the potential shifts in their flower phenology due to competition or reproductive processes. Their impacts on long-term dynamics are beyond the scope of this paper and could be examined in future studies. The model fits to the data are provided in Fig. S5.

We used the per-pot parameters of flowering phenology estimated in model S6 ($fpeak_{i,p}$, $m_{i,p}$ and $\sigma_{i.p}$), to calculate the expected total flower production by species $i$ in each pot $p$ over the growing season. This was achieved by summing up the expected flower production per day:

$$\text{Expected per-pot total number of flowers}_{i,p} = \sum_{d=d_{min}}^{d_{max}} fpeak_{i,p} \exp\left(-\frac{\left(d - m_{i,p}\right)^2}{2\sigma_{i,p}^{\;2}}\right). \quad (S7)$$

$d_{min}$ and $d_{max}$ respectively correspond to the start and end date of the flowering phenology (1 June and 1 November for both species).

Then, we parameterized the competition model (Eq. S2), by fitting the model to the relationship between the expected per-pot total number of flowers produced over the growing season (Eq. S7) and the densities of conspecific and heterospecific individuals within a pot (denoted as $n_{i,p}$ and $n_{j,p}$). We used a hierarchical nonlinear regression model with a negative binomial error distribution. Models with a negative binomial error distribution had better predictive performance than models with a Poisson error distribution according to leave-one-out cross validation (results not shown).

$$\begin{gathered}\text{Expected per-pot total number of flowers}_{i,p} \sim \text{NB}\left(n_{i,p}\, \lambda_{i,source[p]} \exp\left(-\alpha_{ii} n_{i,p} - \alpha_{ij} n_{j,p}\right), \phi\right), \\ \lambda_{i,source[p]} = \exp(\alpha_{ii})\, \lambda'_{i,source[p]} \\ \lambda'_{i,source[p]} \sim \text{N}\left(\lambda'_i, \sigma_{\lambda,i}^{\;2}\right). \end{gathered} \quad (S8)$$

In the estimation, we converted the maximum flower production of species *i*, ($\lambda_i$ in Eq. S2) to $\exp(\alpha_{ii}) \times \lambda'_i$. The rescaled maximum flower production $\lambda'_i$ corresponds to the total number of flowers produced by an individual when it was planted alone (i.e., $n_{i,p} = 1$). This conversion allows us to implement our best knowledge into the Bayesian prior—that Cc and Ccfc individuals produce 2000–4000 seeds over the growing season when grown alone (Katsuhara and Ushimaru 2019). Furthermore, to account for the possibility that individuals of different transplanting classes differ in their lifetime flower productivity, we considered that $\lambda'_i$ differs across the transplanting classes (three classes: early, middle, and late, see details for *2. Competition experiment*), which were assumed to follow a normal distribution N [$\lambda_i$, $\sigma_{\lambda,i}$]. Note that the per-capita flower production, $\lambda_{i,source} \times \exp(-\alpha_{ii}\, n_{i,p} - \alpha_{ij} n_{j,p})$ was multiplied by the number of individuals of species *i* within a pot *p*, $n_{i,p}$, to convert it into pot-level flower production. We used an informative prior for $\lambda'_i$: N [2000, 2000], and vague priors for $\alpha_{ii}$ and $\alpha_{ij}$: N [0, 1], with both restricted to be positive. For the overdispersion parameter in the negative binomial distribution, $\phi$, we used the default prior of the *brm* function: Inv-Gamma [0.4, 0.3]. The parameter estimation procedure was the same in the estimation of phenology curve parameters. The model fits to the data are shown in Fig. S2.

Finally, we parameterized Eq. S3 that describes reproductive interference from the pollination experiment. We fitted a generalized linear mixed-effects model with a binomial error distribution and logit link function to the observed relationship between the number of seeds produced by each flower ($k$) of species $i$ in pot $p$ and $f_{g,d}$, the frequency of conspecific flowers (including both perfect and staminate) within greenhouse ($g$) on the day of marking ($d$):

$$\begin{aligned} \text{Seed number}_k &\sim \text{Binom}\left(4, \frac{\exp(\eta_k)}{1+\exp(\eta_k)}\right), \\ \eta_k &= a_{i,p} + b_i c_g f_{g,d}, \\ a_{i,p} &\sim \text{N}\left(a_i, {\sigma_{a,i}}^2\right), \\ c_g &= \begin{cases} 0 & \text{(greenhouse without bumblebees)} \\ 1 & \text{(greenhouse with bumblebees)} \end{cases}. \end{aligned} \tag{S9}$$

To account for the repeated sampling from the same experimental pots, we included a pot-level random intercept $a_{i,p}$ that follows a normal distribution. To improve the estimation of the random intercept, we used the data from the bumblebee-free greenhouse, in addition to data from the two greenhouses with pollinators. This was achieved by multiplying $b_i$ (the slope representing the conspecific frequency effect) by $c_g$, an indicator variable for pollinator presence ($c_g = 1$ in the greenhouse with bumblebees and $c_g = 0$ otherwise), reflecting that reproductive interference does not occur in the absence of pollinators. Consequently, data without pollinators informed only the random intercept, whereas the slope was estimated solely from data collected in the presence of pollinators. For the parameters $a_i$ and $b_i$, we adopted non-informative priors (Normal [0, 100]).

In all parameter estimations, the model convergence was checked by confirming that R-hat values < 1.01 and by visually inspecting trace plots.

#### *4.2 Prior sensitivity of candidate models*

In the abovementioned parameter estimation, we made best use of our knowledge on the system by specifying the priors in Bayesian inference. For several parameters (all phenology parameters and the rescaled intrinsic flower production), we had biological knowledge reasonable enough to set reliable priors. However, for other parameters, particularly the species-specific competition parameters $\alpha_{ii}$ and $\alpha_{ij}$, we did not have empirical "guesses" and adopted vague priors. Here we check the sensitivity of the parameter estimation to the choice of priors, from which we concluded that Ricker competition function was better suited to our specific system than another candidate function.

Our initial candidate mathematical expressions of competitive dynamics were Ricker and Beverton-Holt (BH) functions:

$$TotalFlower_{i,y} = \begin{cases} \lambda_i \exp\left(-\alpha_{ii}N_{i,y} - \alpha_{ij}N_{j,y}\right) \ (Ricker) \\ \dfrac{\lambda_i}{1 + \alpha_{ii}N_{i,y} + \alpha_{ij}N_{j,y}} \quad (BH) \end{cases} . \qquad (S10)$$

These are among the most common phenological functions of density-dependent competition of plants (Stouffer 2022) and are often reported to be better fitted than other derivatives to observed relationships between plant fecundity and neighbor density (e.g., Levine and HilleRisLambers, 2007; Germain et al., 2016).

We statistically fitted BH and Ricker functions to the observed relationship between the expected per-pot total flower production and the densities of conspecific and heterospecific individuals (see model S8). For the two models, we changed the priors of $\alpha_{ii}$ and $\alpha_{ij}$ from N [0, 1] (our "default") to N [0, 0.1] or N [0, 10] with restricting the values to be positive in all cases and compared the posterior distributions. We found that the posterior distributions of $\alpha_{ii}$ and $\alpha_{ij}$ were substantially influenced by the priors in the BH model, such that increasing the prior standard deviation inflated the posterior estimates (Fig. SM1). In contrast, the posteriors were relatively robust under the Ricker model, except that the posteriors shrank slightly towards zero when a narrower prior (N [0, 0.1]) was used. These contrasting behaviors suggest that competition affecting flower production among these *Commelina* species is better described by the Ricker function, rather than by the BH, for which we used the Ricker function throughout this study. This choice is also biologically reasonable. The BH function is generally considered to represent contest-like competition (winner takes all), whereas the Ricker represents scramble-like competition (everyone shares the loss) (Brännström & Sumpter, 2005). Unlike many plant systems, where competition is dominated by asymmetric vertical competition for light (and thus contest-like, Weiner, 1990), *Commelina* species form dense, low-growing stands that primarily spread horizontally. Competition in this system therefore seems to involve below-ground resources or space rather than light, making scramble competition—and thus the Ricker function—a more plausible representation.

Figure SM1

Sensitivity of the posterior distribution (95% CI) of the parameters in the competition functions (Beverton-Holt [left] and Ricker [right]) to the choice of different priors for αs (N [0, 0.1], N [0, 1], and N [0, 10]). $\alpha_{cc}$ and $\alpha_{ccfc}$ correspond to the competitive effects *from* Cc and Ccfc. Competitive effects *on* Cc and Ccfc are represented with different colors. Note log scales in the y-axis.

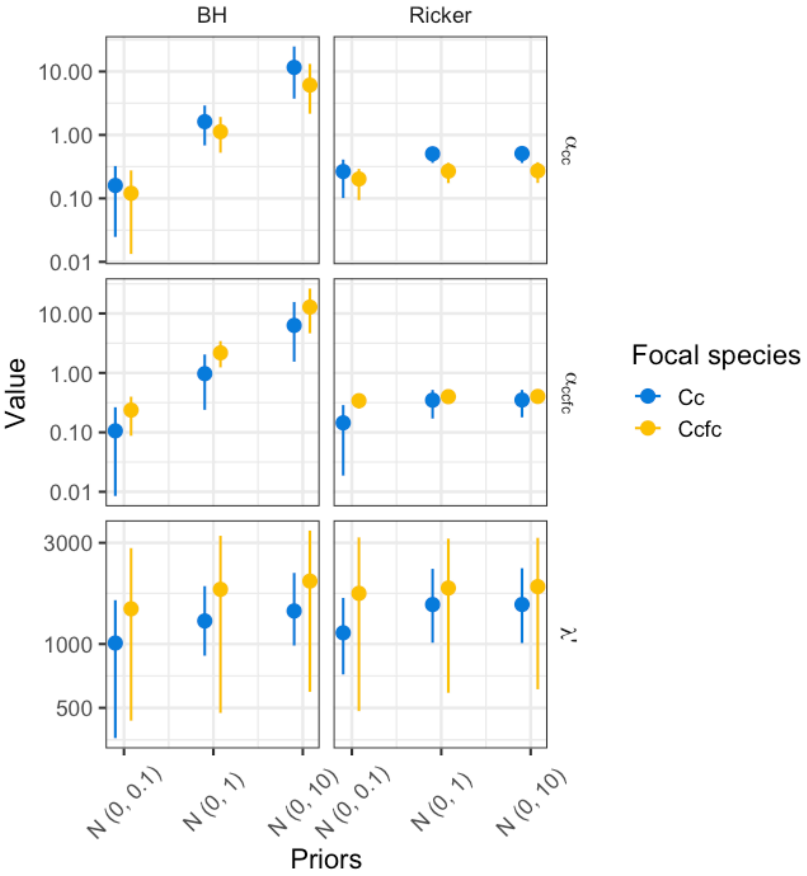


*4.3* Seed germination and survival probabilities

We determined seed germination and mortality rates of *Commelina communis* [:Cc] and *C. c. f. ciliate* [Ccfc], based on previous studies. For these parameters of Cc, we referred to three studies that conducted seed germination experiments and measured the germination and mortality rates (Watanabe and Hirokawa 1971; Suzuki 2003; 2004). Watanabe and Hirokawa (1971) buried seeds of Cc into soil in the same year as harvest and dug them out one year, one year and a half, two years, two years and a half, and four years and a half later. The seeds were then subjected to germination trials in petri dishes in the lab, and the number of germinated and apparently viable seeds were counted. Suzuki (2003) also buried seeds of Cc into soil, dug them out every 2 months up to two years, and conducted germination trials in the field condition and checked seed viability by visual inspection of embryo and endosperm. The germination and survival rates were retrieved from the figures in the papers using the PlotDizitizer (https://plotdigitizer.com/).

For each temporal trajectory of the proportion of viable seeds, we fitted linear regressions with normal distribution, with the response variable being the log-transformed proportion of viable seeds, the explanatory variable the years after burying, and the intercept fixed as log 100. Here, the slopes corresponded to the annual mortality rate of seeds. We also calculated the proportion of germinated seeds among viable ones for each observation in each study, which we interpreted as the germination rate of seeds. The seed survival rate was estimated as 0.734 in Watanabe and Hirokawa (1971) and 0.850 in Suzuki (2003), whereas seed germination rates were 0.807 and 0.795 (Fig. SM2). We therefore decided to use in our model $s_i$=0.8 and $g_i$=0.8 for Cc. Since we could not find any studies measuring germination or survival rates of Ccfc, we assumed the seed germination and survival rates do not differ between these two congeners and used the same values as Cc for Ccfc ($s_i$=0.8 and $g_i$=0.8). In this sense, our results might under- or overestimate their fitness difference arising from the possible inequity of seed survival.

Figure SM2

Temporal trajectories of survival rates (filled circles) and germination rates (proportions of germinated seeds among viable ones, blank circles) of Cc seeds measured in Watanabe and Hirokawa (1971) and Suzuki (2003). Solid lines represent the fit of linear regression between survival rates and year after burying (log [survival rate] = log 100 + log $s_i$ × year, which is equivalent to [survival rate] = 100 × $s_i^{\text{year}}$). The slope estimates of the regression models (i.e., log $s_i$) were –0.311 in Watanabe and Hirokawa (1971) and –0.162 in Suzuki (2003). Broken lines represent the mean of the germination rates calculated separately for the two datasets (0.807 and 0.795).

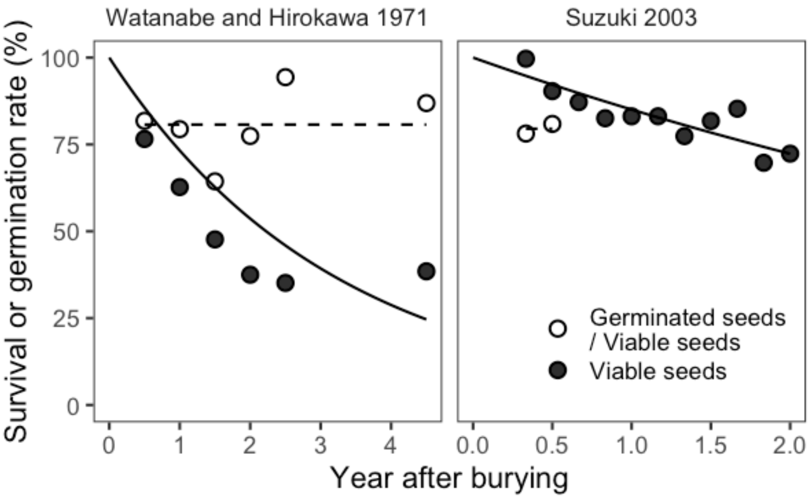
Watanabe and Hirokawa 1971
Suzuki 2003
Survival or germination rate (%)
100
75
50
25
0
0
1
2
3
4
0.0
0.5
1.0
1.5
2.0
Year after burying
Germinated seeds
/ Viable seeds
Viable seeds

### *5. Community dynamics simulations*

*5.1 Coexistence condition without reproductive interference*

When simulating community dynamics using our model, we first considered a case when ecological competition is the only driver of the community dynamics. This mirrors the practice of common coexistence studies that ignore any complexity arising from reproductive interactions (i.e., reproductive interference). To do so, we fixed the number of seeds produced by a flower of each species as its maximum according to our regression model (when the conspecific frequency is one in Fig. 2 in the main text): $MaxSeed_i$ = 1.99 for Cc and 0.521 for Ccfc.

Then, the community dynamics described by Eq. S1 reduce to

$$N_{i,y+1} = (1-g_i)s_iN_{i,y} + g_iN_{i,y}\sum_{d=d_{min}}^{d_{max}} Flower_{i,d,y}p_iMaxSeed_i$$

$$= (1-g_i)s_iN_{i,y} + g_iN_{i,y}p_iMaxSeed_i\sum_{d=d_{min}}^{d_{max}} Flower_{i,d,y}$$

$$= (1-g_i)s_iN_{i,y} + g_iN_{i,y}p_iMaxSeed_iTotalFlower_{i,y}$$

$$= (1-g_i)s_iN_{i,y} + g_iN_{i,y}p_iMaxSeed_i\lambda_i\exp\left(-\alpha_{ii}g_iN_{i,y} - \alpha_{ij}g_jN_{j,y}\right).$$

This is the same form as the conventional model for annual plant competition with Ricker competition, where the coexistence condition is readily available. In this specific model, coexistence is predicted if

$$\sqrt{\frac{\alpha_{ii}\alpha_{jj}}{\alpha_{ij}\alpha_{ji}}} > \sqrt{\frac{\alpha_{ji}\alpha_{jj}}{\alpha_{ii}\alpha_{ij}}}\frac{log\left(\frac{g_i}{1-s_i(1-g_i)}p_iMaxSeed_i\lambda_i\right)}{log\left(\frac{g_j}{1-s_j(1-g_j)}p_jMaxSeed_i\lambda_j\right)} > \sqrt{\frac{\alpha_{ij}\alpha_{ji}}{\alpha_{ii}\alpha_{jj}}}.$$

Conventionally, the left-hand side of this inequity is called as the niche difference, and the right-hand side is its inverse. The middle-hand is the fitness difference, determined by a ratio of the product of sensitivity to competition, survival of seeds in seed bank weighted by germination rates, and maximum seed productivity.

*5.2 Classification and examination of simulated dynamics*

From the posterior draws of the Bayesian regression models, we retrieved 1000 sets of the model parameters. Theory of reproductive interference predicts that the community dynamics would be contingent on initial frequencies as it induces positive frequence-dependence in population growth. To account for this possibility, we started the simulations from three combinations of the seed number of Cc and Ccfc: 49:1; 25:25: and 1:49 for each parameter set. The simulations were run for 100 years. For each of the 3000 simulated dynamics (1000 parameter sets × 3 initial conditions), we classified its *competitive outcome* based on the frequency in the last year: Exclusion of Ccfc (Cc frequency > 0.99); Exclusion of Cc (Cc frequency < 0.01); and Coexistence (otherwise). Then, based on the correspondence of the competitive outcome between the three dynamics with the same parameter set but with different initial conditions, we assigned each parameter set a *dynamical outcome*. For example, if all the three simulations starting from different frequency converged to coexistence, we assigned the parameter set *global* coexistence. When some of the three simulations result in exclusion of Cc and others in exclusion of Ccfc, we assigned *contingent* exclusion (Cc wins or Ccfc wins). In our simulations, six dynamical outcomes were observed among the 1000 parameter sets: three global outcomes (global coexistence, global exclusion of Cc, and global exclusion of Ccfc) and three contingent outcomes (exclusion of Ccfc or coexistence, exclusion of Cc or coexistence, and exclusion of Cc or Ccfc) (see Fig. 4 in the main text).

During the simulations, we noticed that 137 out of the 3000 runs entered an unrealistic state in which flower production by both species declined to nearly zero. This resulted in an ill-defined flower frequency (i.e., zero divided by zero), and, consequently, an ill-defined value of frequency-dependent seed production. This occurred because population overcompensation happened when the abundance of Cc became extremely high, which caused $\exp(-\alpha_{ii}\, n_{i,p} - \alpha_{ij}\, n_{j,p})$ in the Ricker competition function on flower numbers to approach zero (beyond computation). Because Cc was numerically dominant in all runs exhibiting this behavior, we classified these cases as exclusion of Ccfc.

*5.3 Varying degrees of selfing rates and phenology partitioning*

To quantitatively examine the extent to which “selfing rates” (the intercepts in Fig. 3a in the main text) mediate the contingency of community dynamics, we systematically varied the parameter $a_i$ in Eq. S4 to make the intercept range from nearly zero (0.001) to each species’ estimated maximum number of seed production (values when conspecific frequency equals one in the regressions shown in Fig. 3a), while the value when conspecific frequency equals one is fixed (Fig. SM3a). These two extreme cases correspond to complete reproductive interference (individuals cannot produce seeds when rare) and no reproductive interference (conspecific frequency does not affect seed production), respectively. Specifically, we manipulated $a_i$ and $b_i$, by introducing an increment parameter $x$ (ranges from –1 to 1) as:

$$a_i = \begin{cases} log\left(\dfrac{MinSeed_i + (MinSeed_i - 0.001)\ x}{4 - MinSeed_i + (MinSeed_i - 0.001)\ x}\right) & \text{when } x < 0 \\ log\left(\dfrac{MinSeed_i + (MaxSeed_i - MinSeed_i)\ x}{4 - MinSeed_i + (MaxSeed_i - MinSeed_i)\ x}\right) & \text{when } x \geq 0 \end{cases},$$

$$b_i = log\left(\frac{MaxSeed_i}{4 - MaxSeed_i}\right) - a_i.$$

Here $MinSeed_i$ represents the estimated minimum value of the predicted seed production per flower (the regression intercept in Fig. 3a) and $MaxSeed_i$ is the maximum seed production predicted when conspecific frequency equals one. This allows us to modify the minimum seed production while keeping the maximum seed production constant (Fig. SM3a).

We also manipulated the interspecific phenology segregation by altering the flowering peak ($m_i$ in Eq. S3) of Cc and Ccfc (Fig. SM3b), to quantify the role of phenology segregation as well. Specifically, we delayed and advanced the peak of Cc and Ccfc by 1-day increments ($d$: ranging from –30 to 7). Because the estimated peak of Cc was around 14 days earlier than that of Ccfc ($m_i$ =1.83 for Cc; 16.2 for Ccfc), a seven-days shift ($d$=7) results in almost identical peak (hence the smallest phenology segregation, Fig. SM3b). For the greatest segregation, we adopted $d$= –30 (Fig. SM3b). We notice that solely reducing the differences in phenology peaks does not lead to perfect phenology overlap as the curve width ($\sigma_i$ in Eq. S3) remains unaltered. We did not manipulate this parameter for simplification purposes; therefore, these species still partition their flowering phenology to a small extent even in our greatest phenology overlap case ($d$=7).

With all the combinations of varying $x$ and $d$, we ran the community dynamics simulations as described above, except that we fixed the other parameters as the median of their posterior draws for the sake of computational time. For each combination of $x$ and $d$, we started the dynamics from three different conditions (frequencies of Cc) and assigned a dynamic outcome (either *global* or *contingent* ones) based on the correspondence of the three dynamics' competitive outcome.

Figure SM3

(a) We manipulated the "selfing rates" of the *Commelina* species by systematically varying the intercept ($a_i$) of the regression between the number of seeds produced per flower and conspecific frequency. These curves correspond to a range of reproductive interference scenarios from complete reproductive interference (almost zero intercepts, $x = -1$) to none (flat curves, $x = 1$). (b) As a comparison, we also varied the interspecific phenology segregation by delaying and advancing the flowering peak of Cc and Ccfc by 1-day increments. The panels represent three example scenarios: the greatest phenology partitioning ($d = -30$, left); the observed phenology difference ($d = 0$, middle); and the greatest phenology overlap ($d = 7$, right).

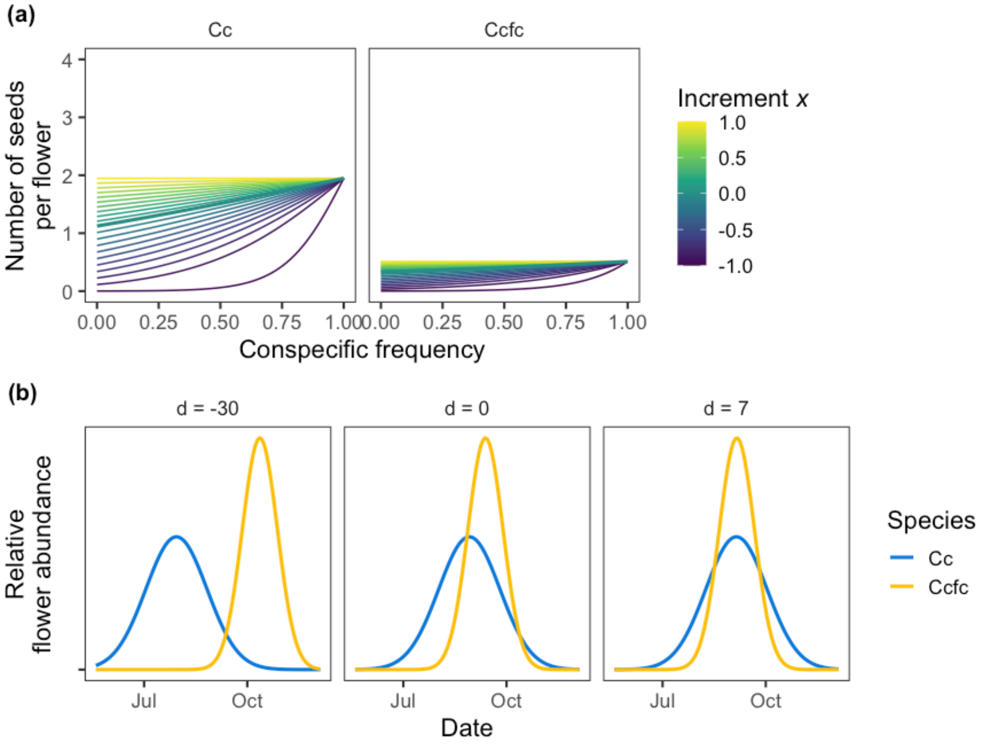

### Supplementary materials

Table. S1

Number of replicates for each combination of the number of Cc and Ccfc individuals in an experimental pot.

| Number of individuals in an experimental pot | | | Number of replicates (for each transplanting class) | | | |
|---|---|---|---|---|---|---|
| Cc + Ccfc | Cc | Ccfc | Total | Early | Middle | Late |
| 1 | 1 | 0 | 16 | 4 | 6 | 6 |
| 1 | 0 | 1 | 18 | 6 | 6 | 6 |
| 2 | 2 | 0 | 8 | 6 | 2 | 0 |
| 2 | 1 | 1 | 10 | 8 | 2 | 0 |
| 2 | 0 | 2 | 8 | 6 | 2 | 0 |
| 3 | 3 | 0 | 4 | 2 | 2 | 0 |
| 3 | 2 | 1 | 2 | 0 | 2 | 0 |
| 3 | 1 | 2 | 6 | 4 | 2 | 0 |
| 3 | 0 | 3 | 2 | 0 | 2 | 0 |
| 4 | 0 | 4 | 2 | 2 | 0 | 0 |
| 5 | 5 | 0 | 6 | 0 | 6 | 0 |
| 5 | 4 | 1 | 4 | 0 | 2 | 2 |
| 5 | 2 | 3 | 2 | 0 | 2 | 0 |
| 5 | 1 | 4 | 4 | 0 | 2 | 2 |
| 5 | 0 | 5 | 6 | 2 | 4 | 0 |

Table. S2

Summary of the mathematical model used to project long-term community dynamics under ecological competition and reproductive interference: (a) Model equations, (b) model variables, and (c) model parameters.

**(a) Model equations**

| Equation | Description |
|---|---|
| Eq. S1: $N_{i,y+1} = (1 - g_i)s_i N_{i,y} + g_i N_{i,y} \sum_{d=d_{min}}^{d_{max}} p_i Flower_{i,d,y} Seed_{i,d,y}$ | Interannual dynamics of seed number. |
| Eq. S2: $TotalFlower_{i,y} = \lambda_i \exp\left(-\alpha_{ii} g_i N_{i,y} - \alpha_{ij} g_j N_{j,y}\right)$ | Effect of germinated individuals on lifetime flower production (competition). |
| Eq. S3: $Flower_{i,d,y} = TotalFlower_{i,y} \frac{1}{\sqrt{2\pi{\sigma_i}^2}} \exp\left(-\frac{(d - m_i)^2}{2{\sigma_i}^2}\right)$ | Phenological curve of seasonal flower production. |
| Eq. S4: $Seed_{i,d,y} = 4 \frac{\exp\left(a_i + b_i f_{i,d,y}\right)}{1 + \exp\left(a_i + b_i f_{i,d,y}\right)}$ | Effect of conspecific flower frequency on seed production per flower (reproductive interference). |
| Eq. S5: $f_{i,d,y} = \frac{g_i N_{i,y} Flower_{i,d,y}}{g_i N_{i,y} Flower_{i,d,y} + g_j N_{j,y} Flower_{j,d,y}}$ | Flower frequency |

**(b) Model variables**

| Name | Description |
|---|---|
| $N_{i,y}$ | Number of seeds of species $i$ in year $y$. |
| $TotalFlower_{i,y}$ | Total number of flowers produced by an individual of species $i$ over the growing season in year $y$. |
| $Flower_{i,d,y}$ | Number of flowers produced by an individual of species $i$ on day $d$, in year $y$. |
| $Seed_{i,d,y}$ | Number of seeds produced by a flower of species $i$ blooming on day $d$, in year $y$. |

**(c) Model parameters**

| Name | Description | Data source for estimation |
|---|---|---|
| $\lambda_i$ | Maximum number of lifetime seed production of species $i$. | Competition experiment |
| $\alpha_{ii}$, $\alpha_{ij}$ | Competitive effects of germinated conspecific (species $i$) and heterospecific (species $j$) individuals on flower production of species $i$. | Competition experiment |
| $m_i$ | Peak date of the flowering phenology of species $i$. | Competition experiment |
| $\sigma_i$ | Standard deviation of the flowering phenological curve of species $i$. | Competition experiment |
| $a_i$ | Baseline level of per-flower seed production of species $i$ when the conspecific frequency is zero. | Pollination experiment |
| $b_i$ | Effect of increasing conspecific flower frequency on per-flower seed production of species $i$. | Pollination experiment |

| | | |
|---|---|---|
| $g_i$ | Germination rate of seeds of species $i$. | Previous study |
| $s_i$ | Survival rate of ungerminated seeds of species $i$. | Previous study |
| $p_i$ | Proportion of perfect flowers among all flowers (perfect and staminate) produced by species $i$ | Competition experiment |
| $d_{min}$，$d_{max}$ | Number of seeds produced by a flower of species $i$ blooming on day $d$, in year $y$. | Fixed arbitrary |

Table. S3

Results of a generalized linear model with a binomial error distribution and logit link function relating the proportion of perfect flowers among all flowers (including both perfect and staminate ones) to species identity (categorical: Cc or Ccfc, with Cc as the reference level), the total number of individuals within the same pot (1–5), and transplanting class (categorical: early, middle, or late, with early as the reference level). The estimate, standard errors, and p-values are shown for each explanatory variable.

| | | Estimate | SE | p |
|---|---|---|---|---|
| Intercept | | –0.129 | 0.048 | 0.008 |
| Species: Ccfc | | –0.155 | 0.040 | < 0.001 |
| Total individuals | | –0.006 | 0.013 | 0.651 |
| Transplanting class: | Middle | 0.037 | 0.041 | 0.362 |
| | Late | –0.025 | 0.060 | 0.673 |

Figure. S1

Posterior distributions of the parameters in the competition model. Thick and thin bars respectively represent the 80% and 95% credible intervals, and points are the median.

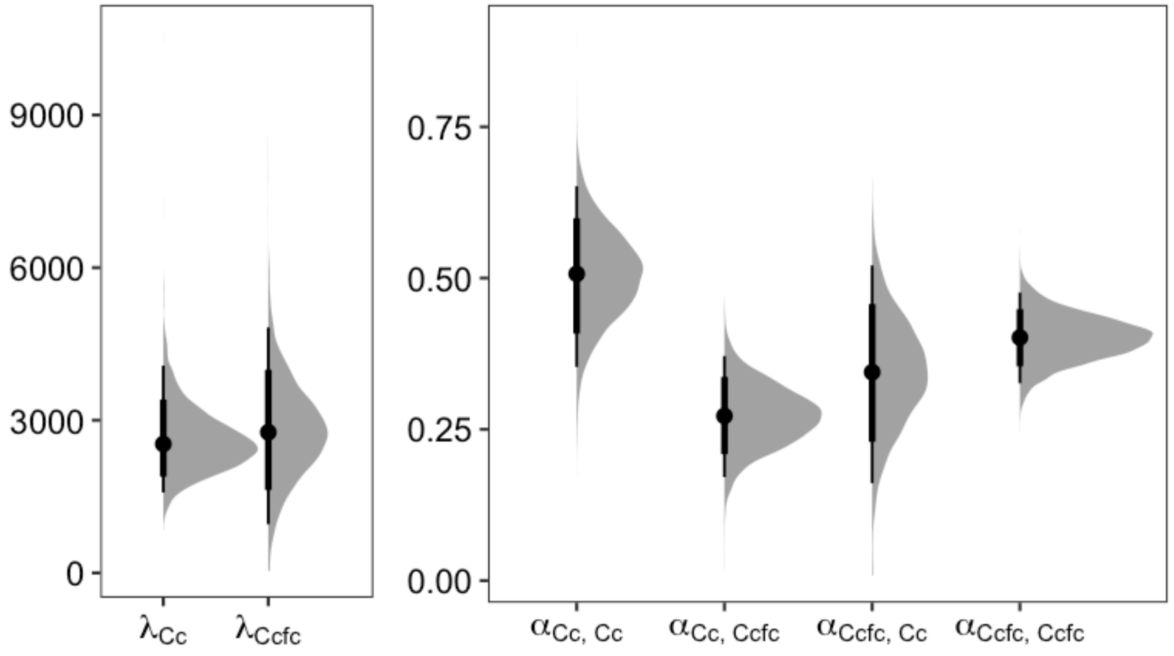

Figure. S2

Relationships between the expected total flower production over the growing season of an individual of (a) Cc and (b) Ccfc and the number of individuals (either Cc [left column] or Ccfc [right column]) within a pot. Points represent pots, with color being their transplanting classes (see more details in *2. Competition Experiment*). Lines and shaded areas indicate the median predictions and 95% credible intervals.

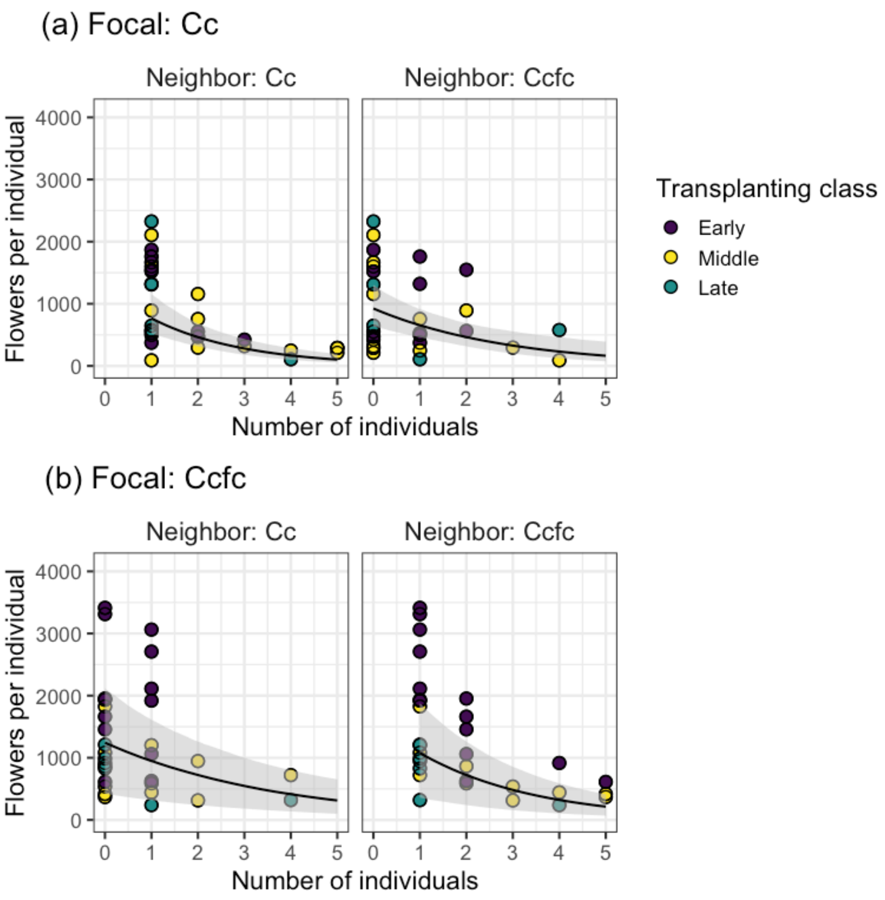

Figure. S3

Comparison of seed production in the emasculation experiment. Because flowers were emasculated (i.e., their stamens were removed) shortly after flower opening and left unpollinated in the bumblebee-free greenhouse, their seed production was possible only through prior selfing. We counted seed production of 87 and 135 flowers of Cc and Ccfc, respectively. Seed production was higher for Cc than Ccfc (the Wilcoxon rank sum test, $p < 0.001$), suggesting a higher rate of prior selfing in Cc. Open points represent individual flowers, and black points indicate species means. The error bars represent ± 2SE.

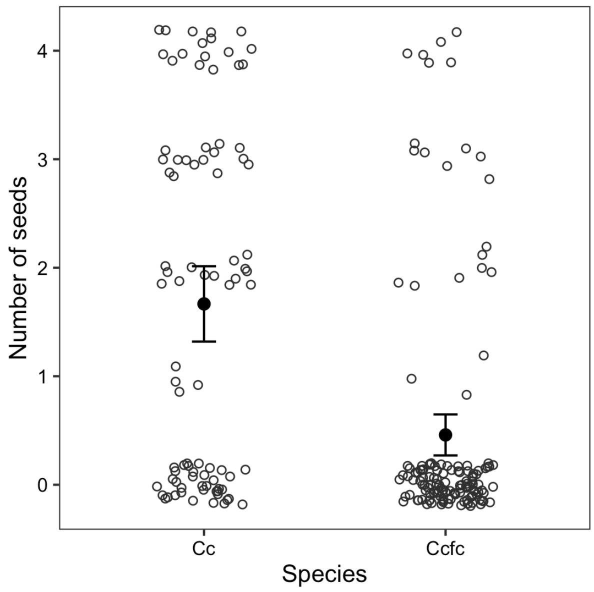

Figure. S4

Comparison of ecological traits of *Commelina communis* (Cc) and *C. c.* f. *ciliata* (Ccfc). P-values calculated from the Wilcoxon rank sum exact test are shown.

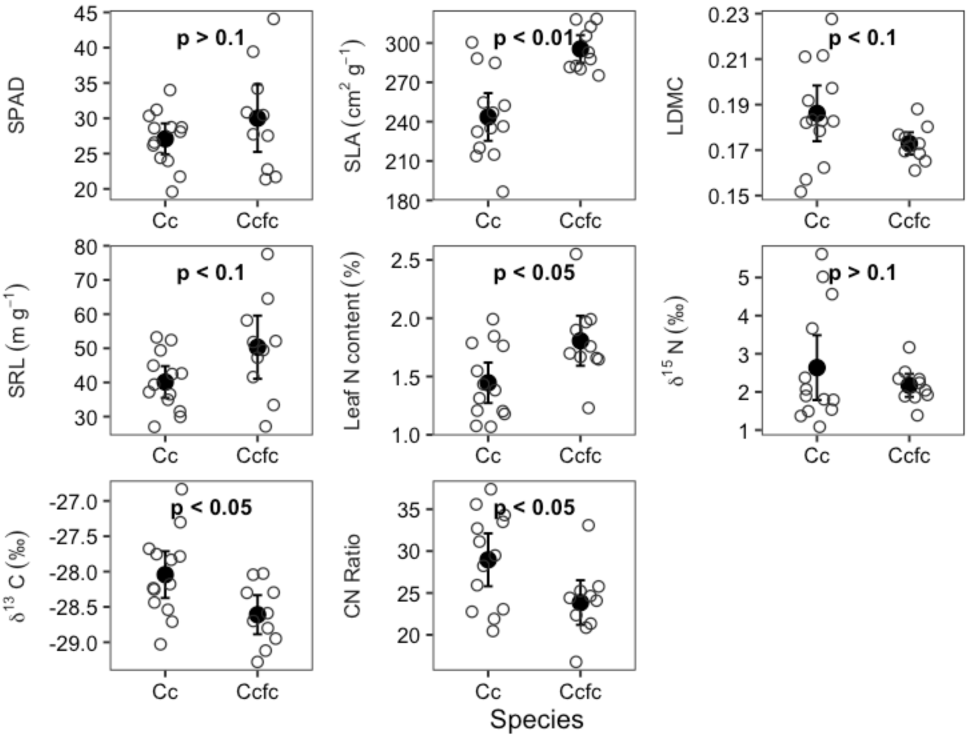

Figure. S5

Observed (open circles) and predicted (lines) seasonal dynamics of flower production of (a) Cc and (b) Ccfc in each pot.

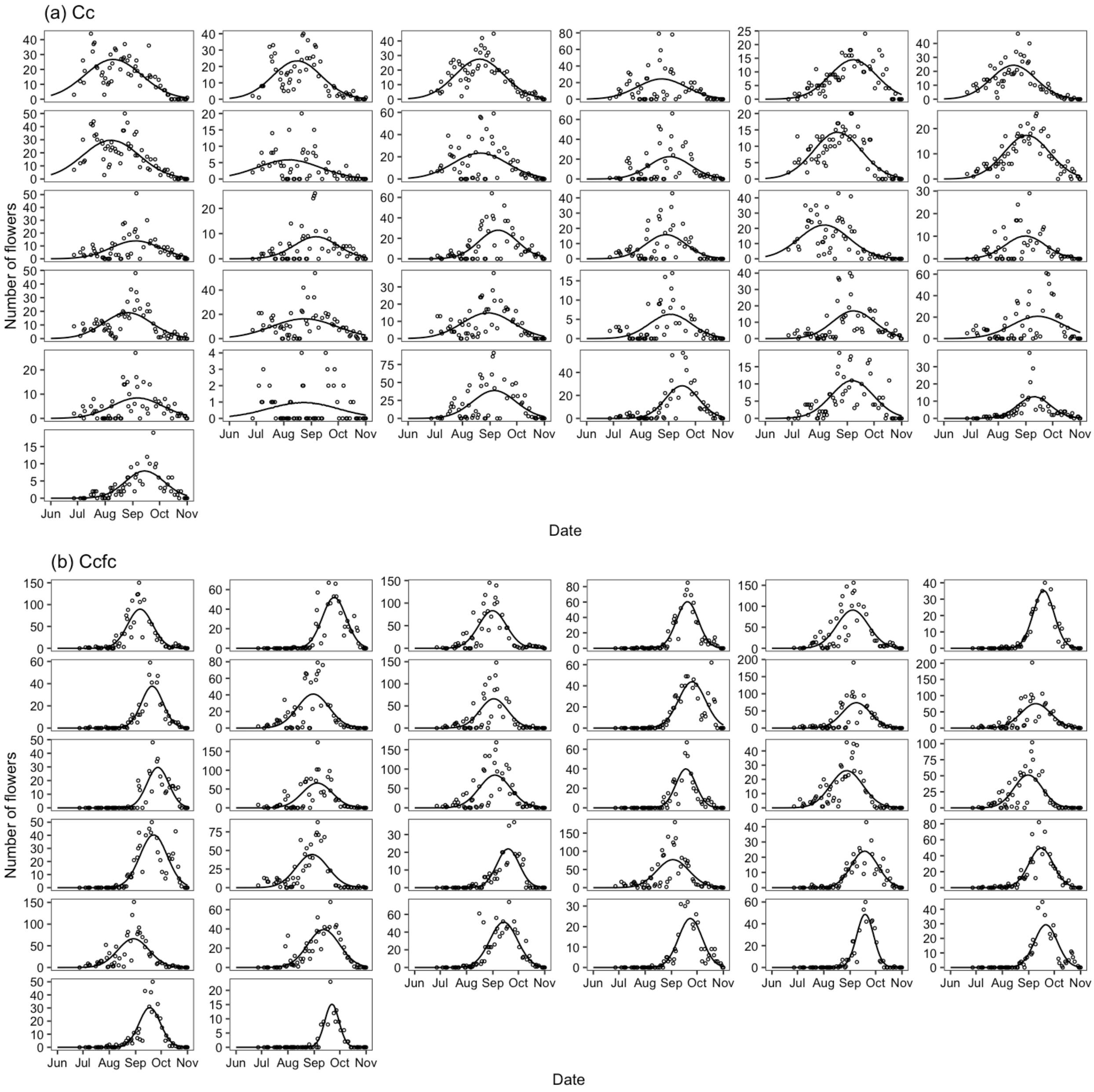
